## Additional File 2 for "Temporal dynamics of gut microbiomes in non-industrialized urban Amazonia"

**Additional file 2: Fig S1**

Nutrient intake across time points as calculated from three-day self-reported dietary records from Belém, Brazil individuals (UN). A) Average açaí consumption (mL) in T1 and T2. B) Average intake of animal protein (g) in T1 and T2. C) Average intake of plant protein (g) in T1 and T2. D) Average intake of fats (g) in T1 and T2. E) Average fiber intake (g) in T1 and T2. F) Average consumption of dairy products (g) in T1 and T2.

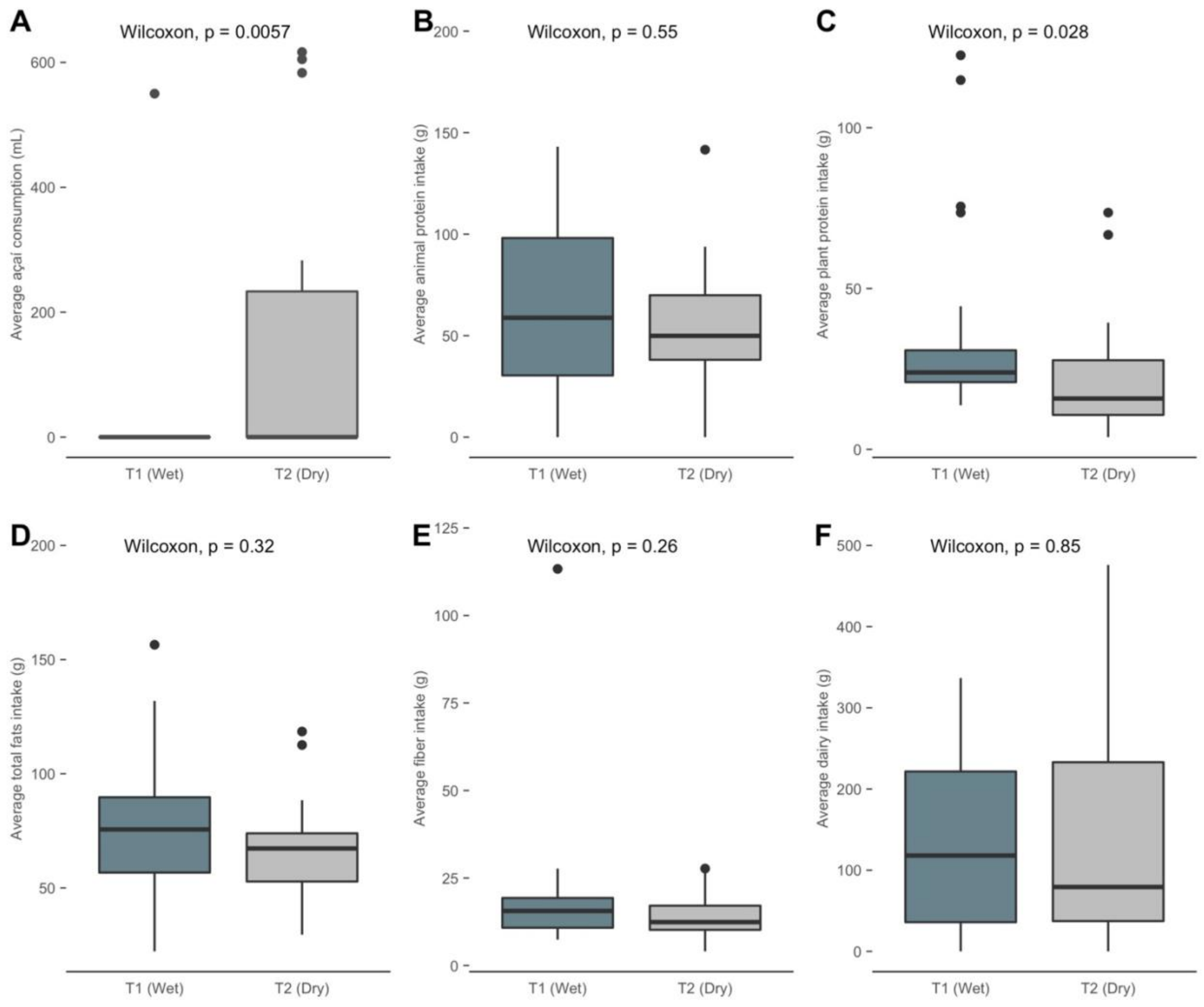

**Additional file 2: Fig S2**

Shannon alpha diversity metrics for United States (UI), Brazil (UN), and Tanzania (RN) individuals. Shapes indicate sample collection times.

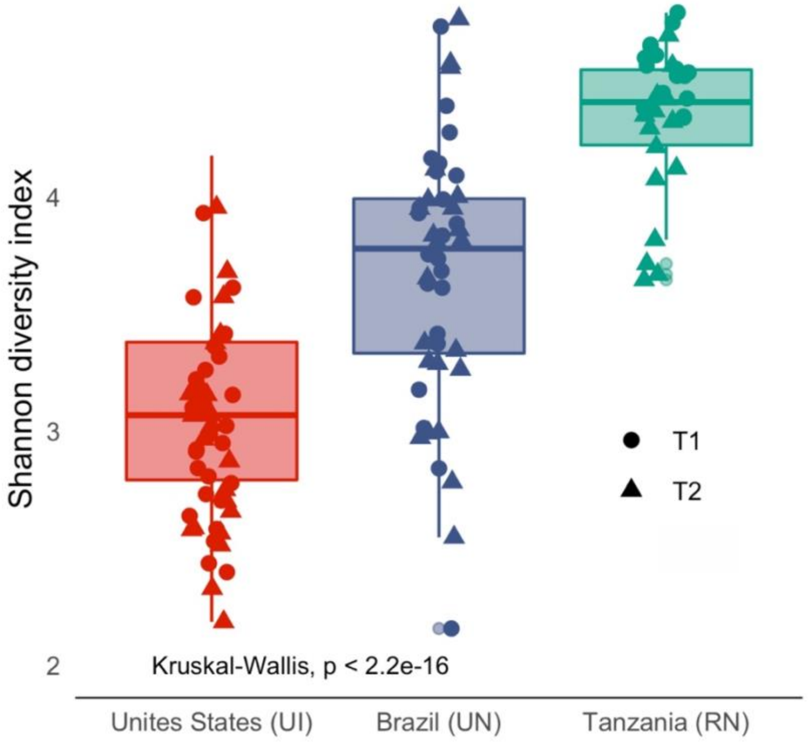

**Additional file 2: Fig S3**

A) Abundance plots of core taxa (genera prevalent in at least 50% of individuals from each location) according to collection time points. B) Abundance plots of level 2 KEGG pathways per population and time points.

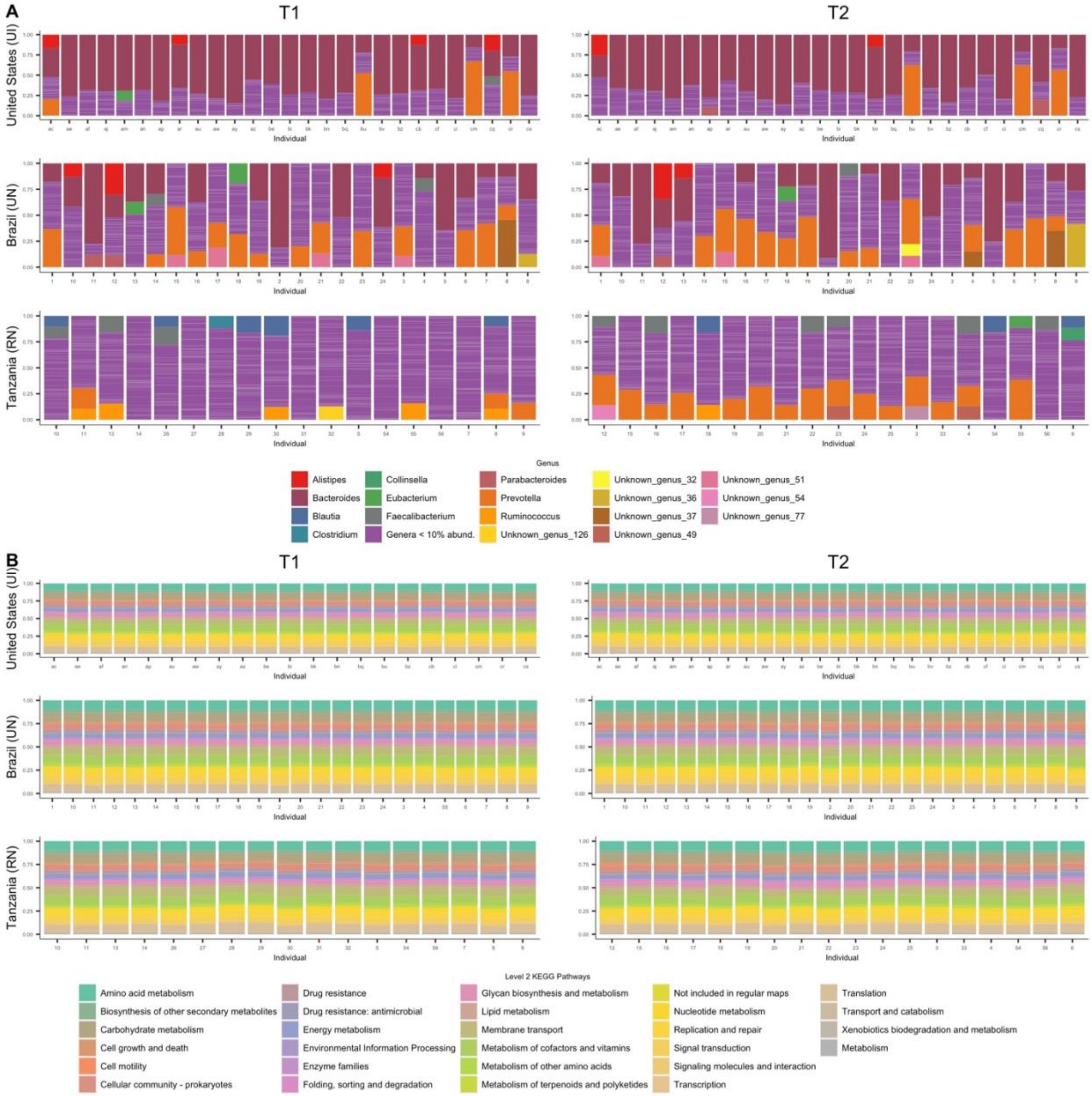

**Additional file 2: Fig S4**

Differential abundance of level 2 KEGG pathways across time points among the Tanzania Hadza hunter-gatherers (RN) according to ANCOM results.

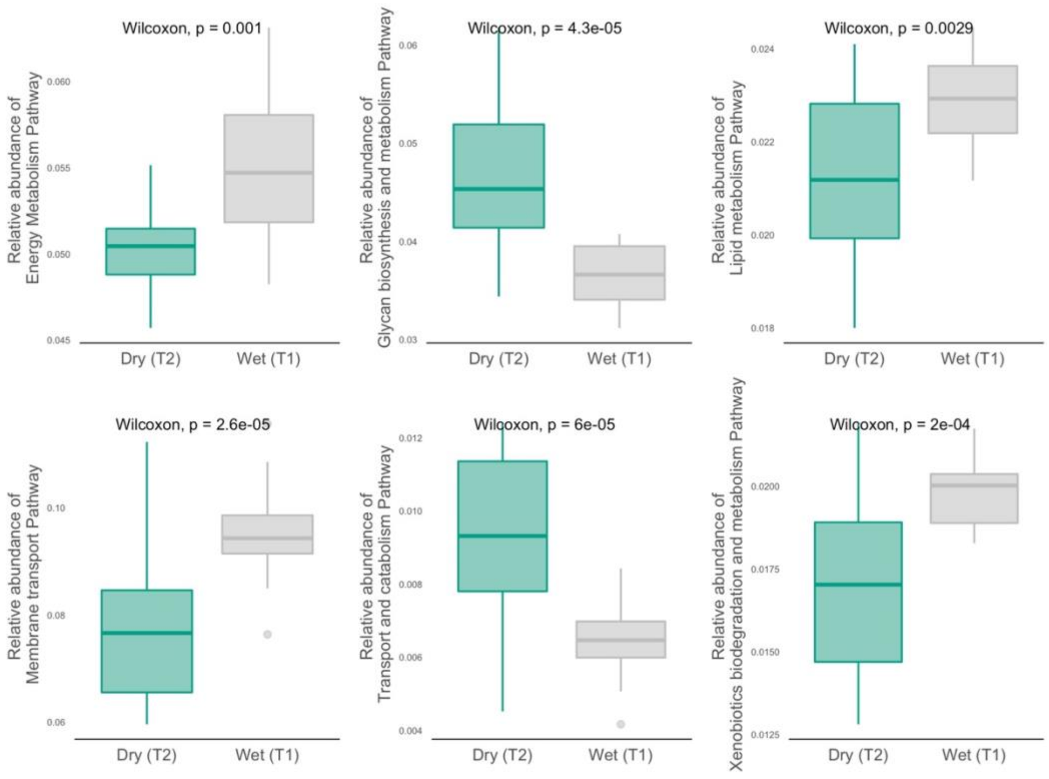

**Additional file 2: Fig S5**

A) Top driver taxa of the four enterotypes attributed to individuals from Brazil (UN), United States (UI), and Tanzania (RN). B) Body mass index values per enterotype classification for individuals from Brazil.

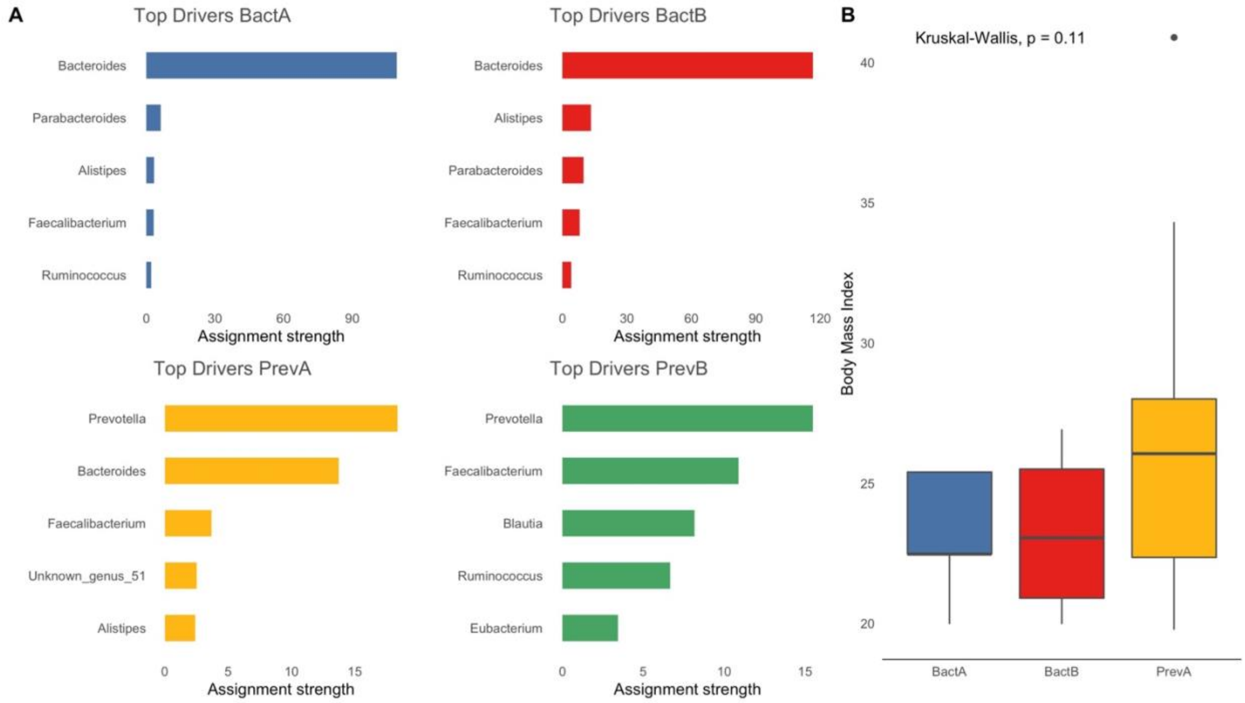

**Additional file 2: Fig S6**

Our approach to investigate strain dynamics across time points. A) InStrain (27) was used to map metagenomic reads from each time point to dereplicated assembled metagenomes of each individual. B) A Consensus SNP is called when the major allele at a given site differs between time points but there are shared alleles between metagenomic reads. A Population SNP is called when no alleles are shared between reads at different time points. C) Demonstration of data filtering for SNP analysis, in which we selected sites with major allele frequencies above the threshold of 0.7 and regarded the remainder as missing data.

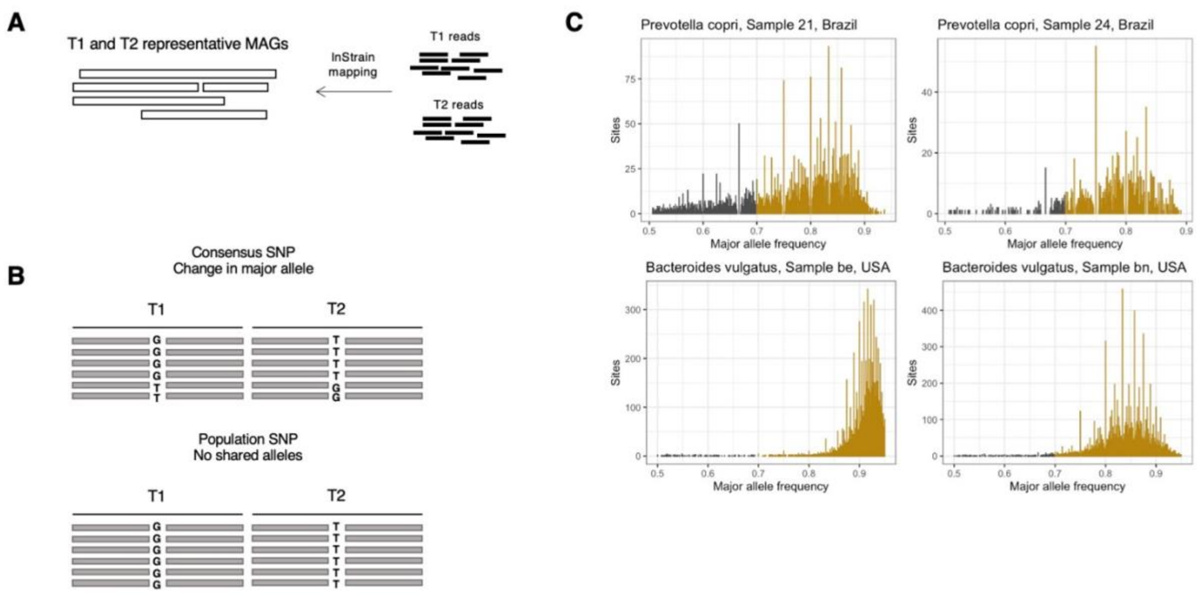

**Additional file 2: Fig S7**

Taxonomic classification of metagenomic reads when using the standard Kraken2 database and when implementing a custom database containing sequences from the Global Microbiome Conservancy (7) isolate library.

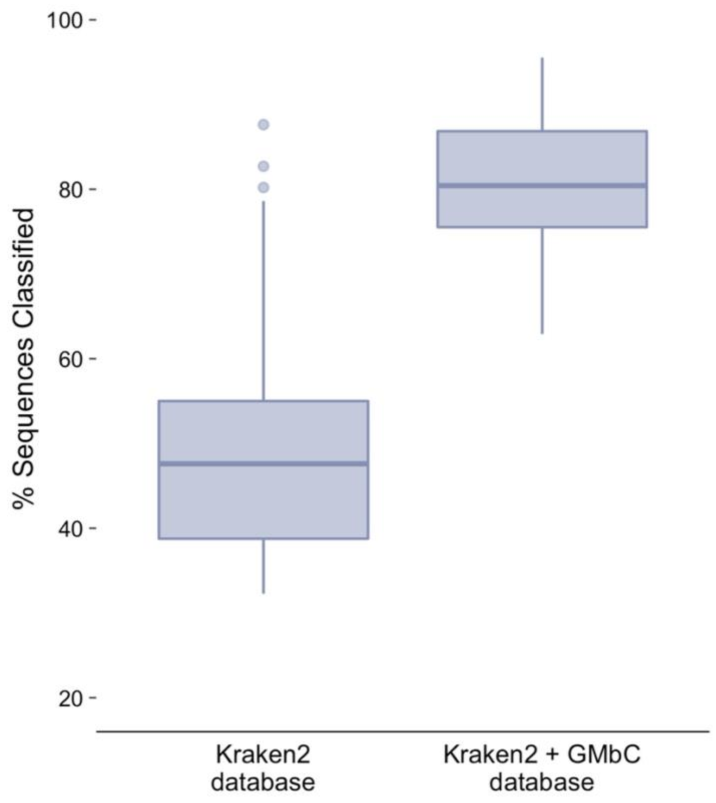
